# Neurocognitive dynamics of translating information from a spatial map into action

**DOI:** 10.1101/2025.10.17.683083

**Authors:** Maud Saulay-Carret, Clément Naveilhan, Xavier Corveleyn, Stephen Ramanoël

## Abstract

How do we translate information from a spatial map to action in our immediate surroundings? Despite the widespread use of various tools for orientation, from paper maps to GPS, this fundamental question remains unanswered in our understanding of human spatial navigation. To investigate this, we implemented a perspective-taking task in immersive virtual reality combined with mobile EEG, aiming to disentangle the neurocognitive processes involved. Thirty-eight young adults were presented with a virtual 2D map in which we manipulated both the perspective shift and the physical angle of rotation required to align with a target, as well as the congruency between these two variables. Behaviourally, angular error during pointing increased slightly and linearly with perspective shift. However, the relationship between rotation angle and accuracy revealed a non-linear pattern, with better performance around the antero-posterior bodily-axis. Regarding congruency, angular error increased for incongruent trials, but only when the perspective-taking angle exceeded 90°. At the neural level, activity in the retrosplenial complex (RSC) revealed a sequential organization with alpha-band modulation during perspective shift, followed by beta-band activity reflecting preparation for the required physical rotation. In addition, incongruency between perspective-taking and physical rotation increased beta activity in the left temporo-parietal junction (lTPJ). Overall, these findings demonstrate the value of immersive virtual environments to investigate the neural correlates of real-world navigation and the complexity of perspective-taking mechanisms.

## Introduction

Among our various spatial navigation abilities, the efficient encoding and use of visuospatial information from maps of the surrounding environment represents an important aspect of orientation skills (He et al., 2022; Keehner et al., 2006; Wolbers & Hegarty, 2010). This complex multi-step process requires first that we mentally orient ourselves in the desired direction to encode the information from the 2D map. Then, this spatial representation must be used in the 3D environment to physically rotate to align with the destination before traveling. Although this ability represents a classic case of spatial orientation, we still lack a clear understanding of how these different operations interact to support efficient navigation, as well as of their neural correlates.

To investigate the neurocognitive processes underlying the integration of different spatial representations, researchers have notably used spatial perspective-taking tasks. These paradigms assess an individual’s ability to adopt a visuospatial perspective different from their own, often employing paper-and-pencil or desktop-based setups such as the Spatial Orientation Task (SOT; Hegarty & Waller, 2004; Kozhevnikov & Hegarty, 2001a) or its revised computerized version, the Perspective Taking/Spatial Orientation Test (Friedman et al., 2020). Analysis of the absolute angular error (*i.e.,* the difference between the direction indicated by the participant and the expected direction) provides information about an individual’s ability to mentally orient to different spatial perspectives. Despite the interest in this field, relatively few experiments have investigated the neural correlates of perspective taking in humans, with considerable heterogeneity in the protocols used (for recent review see Gunia et al., 2021). For example, Ding et al. (2025) recorded scalp electroencephalography (EEG) activity while participants performed a map-based self-localisation task. The authors showed that spatial perspective taking engages a distributed network comprising parietal, frontal, and motor cortices. The study revealed that greater angular disparities were associated with increased P300 amplitudes, along with lateralised motor activity that is interpreted as a marker of embodied motor imagery. In the same way, Gunia et al. (2024) investigated mental rotation shifts between different viewpoints in spatial perspective-taking tasks using intracranial electroencephalography. Their results indicated the involvement of an extensive brain network, including the precuneus, superior parietal lobule, the insula, temporo-parietal junction, lateral prefrontal cortex, and the retrosplenial complex (RSC).

Among this network of brain regions, the RSC represents a central hub for spatial cognition, nestled between cortical and subcortical regions, including the hippocampal, parahippocampal, and visual systems (Alexander et al., 2023; Mitchell et al., 2018; Stacho & Manahan-Vaughan, 2022). It is thought that, through its reciprocal connections with the thalamic and parietal regions, the RSC computes head direction dynamics via the complex integration of visual, vestibular, and proprioceptive information (Cho & Sharp, 2001; Keshavarzi et al., 2023; Sit & Goard, 2023). This central role is reflected in the involvement of the RSC in imagined perspective shifts, when participants anticipate viewpoint changes, as showed by a previous fMRI study (Sulpizio et al., 2013). Further work also proposed that the RSC supports transitions between multiple reference frames (Shine et al., 2016) and integrates different spatial viewpoints, enabling flexible reorientation towards novel perspectives like a neural compass (Alexander et al., 2023; Dhindsa et al., 2014; Gomez et al., 2014; Lu et al., 2025; Marchette et al., 2014).

Recent advances in virtual reality technology and mobile EEG have extended these findings to freely moving humans (Do et al., 2021; Gramann et al., 2021; Griffiths et al., 2024; Lin et al., 2015; Naveilhan et al., 2025). These findings highlighted the importance of slow-frequency brain oscillations in the RSC region (mainly theta-alpha activity, between 2 and 12 Hz), reinforcing the key role of the posterior parietal cortex for perspective taking and navigation in general (Whitlock, 2017; Whitlock et al., 2008). Beyond the mere recording of movement-related activities, several works emphasized the role of body-related cues (*i.e.,* optic flow, motor efference copy, and vestibular inputs) in spatial navigation abilities (Alexander et al., 2023; Stangle et al., 2023; Epstein et al., 2017; Iggena et al., 2023; Moon et al., 2023; Muto et al., 2018). The integration of body-related cues is particularly relevant for perspective-taking, and can either depend on the embodied simulation of one’s own movement or on the anchoring of an external perspective to the body’s orthogonal axis (Kessler & Rutherford, 2010). In line with this view, several studies reported the role of body axes as a stable reference grid around which space can be organised (Iachini, 2024; Mou & McNamara, 2002).

In relation to this growing recognition of the importance of body-related cues in spatial cognition, He et al. (2022) developed and validated a more ecological experimental version of the perspective-taking task (PTT). Their immersive Viewpoint Transformation Task (iVTT) included both perspective-taking and physical rotation phases toward a target presented on a map. Their results suggested that the iVTT engaged similar processes as the computerized version, but also some differences related to the involvement of body-related processes. However, the purely behavioural nature of the task prevented them from delving deeper into the underlying brain dynamics. The combination of this immersive paradigm with mobile neuroimaging may therefore provide a unique opportunity to investigate the neurocognitive processes involved in the integration of different spatial representations, including the physical rotation phase during reorientation.

To this end, we extended the iVTT experimental protocol used by He et al. (2022) and combined it with a mobile EEG approach. Specifically, we finely manipulated the perspective-taking angle and the angle of rotation required to perform the task in order to disentangle the neurocognitive processes involved in spatial perspective taking and the precomputation of physical rotation. We aimed to explore the parameters contributing to the angular error in the task, arguing that the size of the perspective shift may affect the error differently compared to the amount of rotation needed. We also speculated that the congruence between these spatial representations (*i.e.,* is the perspective shift in the same direction as the angle to be executed or not) would affect the task performance, with lower accuracy for incongruent trials reflecting conflict between mental and physical rotations. At the neural level, we expected to identify a specific sequential slow-frequency signature in a cluster of dipoles reconstructed around the RSC, modulated by the congruency between perspective-taking and physical rotation, due to the central role this brain region plays in integrating differential spatial representations.

## Methods

### Participants

40 young adults were involved in the current study. This sample size was determined by *a priori* power analysis based on data from He et al. (2022). We performed a reanalysis of their data from their <u>GitHub</u>, using linear mixed models as we planned to use in the current paper. We fitted a linear mixed-effects model: *lmer(meanAngularError ∼ PerspectiveShift * PointingAngle + (1 | ID))*. From this model, we computed the marginal R², which was then converted to Cohen’s f^2^. Using the *pwr* package (Champely et al., 2020), we estimated that a sample size of 34 participants would be required to achieve 90% power at an alpha level of .05. Participants were compensated €20 for their participation, which lasted approximately two hours. All had normal or corrected-to-normal vision, and no history of neurological or cognitive disorders. The study was approved by the local ethics committee (CERNI-UCA authorization no. 2023-20), and participants gave their informed consent before participation.

Two participants were excluded from all the analyses because of cybersickness as assessed by the motion sickness questionnaire (Kennedy et al., 1993) resulting in a final sample of 38 participants (*M =* 23.63 years; *SE* = 3.59 years; range 18-35 years; 22 female participants). Participants also completed the computerized version of the Perspective Taking/Spatial Orientation Test (Friedman et al., 2020) in order to test their 2D perspective-taking abilities, and compare the performances with the immersive protocol.

### Stimuli and Procedure

Visual stimuli were generated using Unity Engine (Version 2021.3.8f1), and the protocol is freely accessible on the OSF repository of the study. The protocol was adapted from the paradigm provided by He et al. (2022). Participants were immersed in the environment using the HTC VIVE Focus 3 (HTC Corporation) (**Figure 1.B**). This virtual reality headset features two 2.88’ LCD panels with a resolution of 2448 x 2448 pixels per eye (4896 x 2448 pixels combined); a 90 Hz refresh rate, and a field of view up to 120°. Real-time broadcasting was provided by the VIVE Business Streaming Software (HTC Corporation, Version 2.0.8a). The software enabled live broadcasting from the Unity interface using the VIVE Wave XR plugin and the Direct Preview function, using a USB cable. The Unity software was implemented on a Dell Precision 3581 laptop (CPU: i9-13900H 64GB RAM; GPU: NVIDIA RTX 2000 Ada Laptop 8GB vRAM).

**Figure 1.**
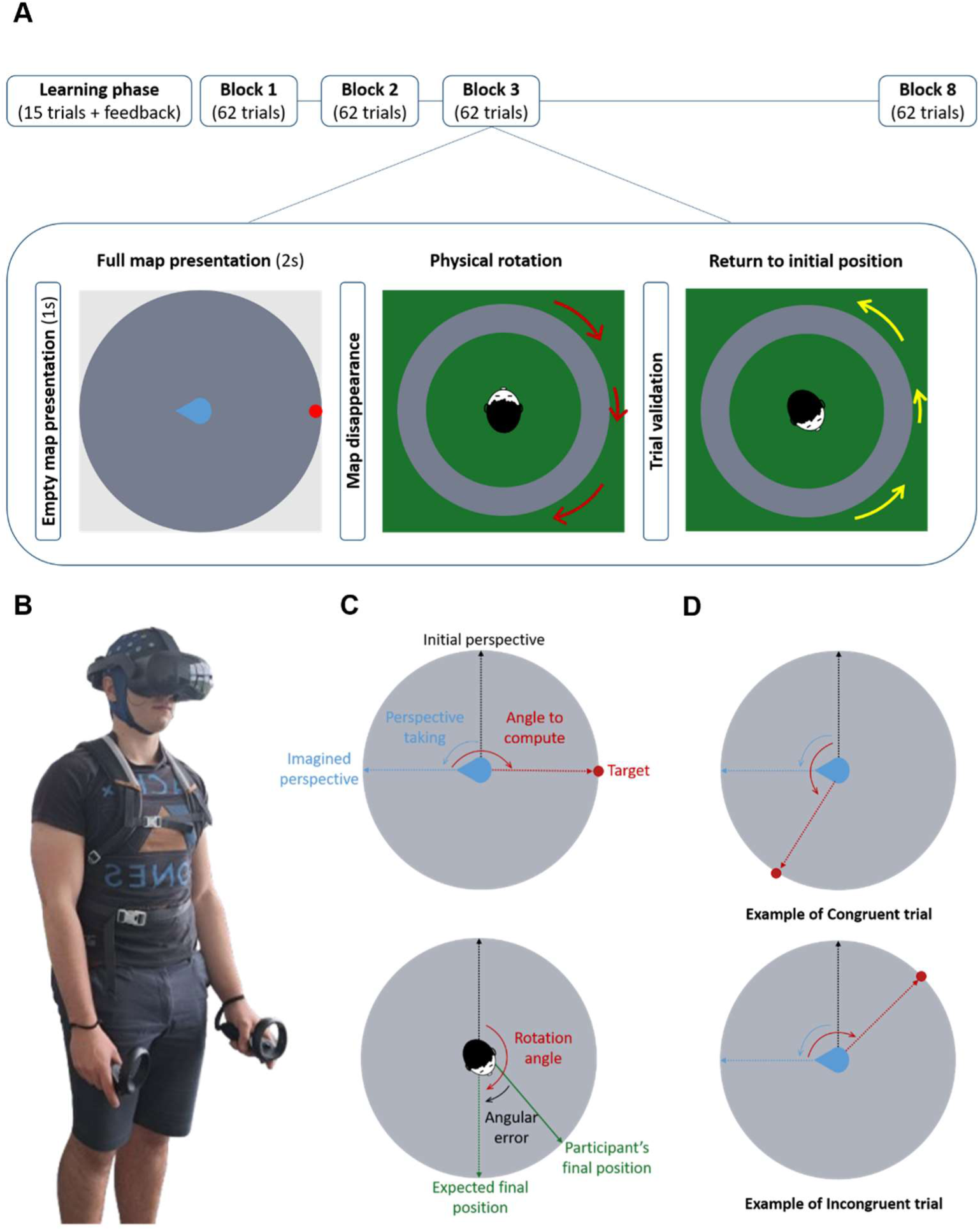
Presentation of the paradigm. **A.** iVTT paradigm. Here, the map illustrates a trial in which participants took an imagined perspective 90° to their left (blue arrow). The target location (red dot) was positioned in the opposite direction, requiring participants to rotate accordingly to align with the goal’s position. Note that the aerial views of the physical rotation and return to initial position are for illustrative purposes only: participants did not have access to this view. **B.** Experimental set-up. **C.** *Perspective-Taking angle* refers to the angle between the *Initial Perspective* (*i.e.*, the participant’s initial physical position) and the *Imagined Perspective*. *Angle to Compute* is the angle between the Imagined Perspective and the *Target*, defining the required movement to align with the pointing direction (and so is equal to the *Rotation Angle*). *Final Position* corresponds to the participant’s position when validating their answer. *Angular Error* is the difference (in degree) between the participant’s final position and the expected final position. **D.** Trial *Congruency* is based on the relationship between the *Perspective-Taking* angle and the *Angle to Compute*. Here, the participant takes an imagined perspective 90° to the left. The figure shows a congruent trial, where the target lies between 0° and 180° after the shift, and an incongruent trial, where it falls outside this range, requiring the participant to mentally “go back” in the opposition direction to locate the correct target position.

Participants were placed in a first-person view in an arena of three metres in radius, surrounded by grey walls and devoid of other visual cues that could be used as landmarks (**Figure 1.A**). A red ring marked the centre of the arena, where participants began the first trial. They first performed a learning phase of 15 trials during which they received feedback and could be guided. After this phase, they proceed to the experimental phase, which consisted of 8 blocks, each containing 62 trials. Once in the centre of the virtual arena, a map of the environment appeared in front of them. Initially, the map was empty (for 1 second), after which a red dot and a blue arrow appeared on it. Participants had to imagine facing in the direction indicated by the arrow (*i.e.,* the perspective taking), adopt this perspective to locate the target point (the red dot) and then, when the map disappeared physically turn toward it (**Figure 1.C**). Once they reached the position they believed to be correct, they validated their response by pressing a button on the right controller. Directional arrows then appeared to guide them back to their initial starting position before proceeding to the next trial.

Participants completed 8 blocks of 62 trials each, for a total of 496 trials and approximately 75 minutes in the virtual environment. They could take as many self-paced breaks as necessary. These 496 trials allowed to explore the full range of possible combinations between 9 perspective angles (from 0 to 360° every 22.5°) and 31 rotation angles (from 0 to 360° every 12°). To achieve a finer coverage, that is, rotation angles measured every 6°, while minimizing participant fatigue, they were divided into two groups. The first group performed tasks with angles spaced at 12° intervals starting in a clockwise direction, while the second group focused on angles spaced at 12° intervals in an anti-clockwise direction. We ensured that there was no difference in task performance across groups (*p* = .640). We also ensured there was no effect of left and right direction for the rotation angle (*F*_(1, 112)_ = 0.019, *p* = .889) and for the perspective taking angle (*F*_(1, 112)_ = 1.497, *p* = .224), and we thus used corrected angles (between 0 and 180 without accounting for direction) for the analyses reported. We distinguished between congruent trials and incongruent trials based on the difference between the direction of the *perspective-taking angle* (blue arrow) and the direction of the *angle to compute* for physical rotation. A trial is congruent if the target location lies between 0° and 180° after the new perspective is imagined; otherwise, if the target is outside this range, the trial is considered as incongruent (**Figure 1.D**).

### EEG Acquisition and Processing

EEG signals were acquired at a sampling rate of 500 Hz using a 64-channel Waveguard original cap equipped with Ag/AgCl electrodes. The electrodes were connected to an eego mylab amplifier from ANT Neuro and digitized at 24-bit resolution. The reference electrode was CPz and the ground electrode was AFz. Impedances were kept below 10k Ω. EEG recordings and stimulus presentations were synchronized using the LabStreamingLayer software Labrecorder (LSL Labrecorder version 1.13; Kothe et al., 2025). EEG data were preprocessed offline using MATLAB (R2021a; The MathWorks Inc.) with custom scripts from the EEGLAB toolbox (version 14.1.0b; Delorme & Makeig, 2004) and the BeMobil pipeline (Klug et al., 2022).

We first downsampled to 250 Hz before automatically and manually removing acquisition segments where the participant was not actively engaged in the task (*e.g.,* at the beginning and at the end of each block or due to any technical problems) or segments containing artefacts. Spectral peaks from line noise and screen refresh rate were then automatically detected and removed using the Zapline-plus algorithm (Klug & Kloosterman, 2022). We identified noisy electrodes using the *clean_rawdata* plugin with default parameters and 20 rejection iterations. On average, 2.18 ± 1.45 electrodes per participant were rejected and reconstructed using spherical interpolation with signals from neighbouring electrodes, followed by re-referencing to the common average. We then applied a transient high-pass filter at 1.5 Hz to the data, as proposed by Klug & Gramann (2021), before performing independent component decomposition using the Adaptive Mixture Independent Component Analysis (AMICA) algorithm (Palmer et al., 2008), which is considered the most effective for EEG data (Delorme et al., 2012). We performed 2000 iterations with 10 rejection iterations using a threshold of 3 standard deviations (Klug et al., 2024) to limit artefacts in the data without applying temporal domain cleaning. Each independent component (IC) was then assigned a dipole source reconstructed using the equivalent dipole model (DipFit ; Oostenveld & Oostendorp, 2002). To classify the components into seven classes, we used the Lite version of the ICLabel algorithm (Pion-Tonachini et al., 2019). After these preprocessing steps, we extracted the IC that was classified as brain with at least 20% probability and residual variance below 15% (Delorme et al. 2012), resulting in an average retention of 20 ± 3.96 components per participant. We then copied this information to the unfiltered cleaned data.

The resulting cleaned signal was filtered with a bandpass filter between 0.3 Hz (with a transition bandwidth of 0.5 Hz, a slope at 0.55 Hz, and an order of 1650) and 80 Hz (with a transition bandwidth of 20 Hz, a slope at 80 Hz, and an order of 42). We then segmented the signal into epochs around the events of interest (*i.e.,* the appearance of the map or the rotation). We then removed epochs containing artefacts greater than 150 µV, resulting in an average of 458.78 ± 71.70 epochs for the map.

### Source reconstruction procedure

After extracting the data, we performed source reconstruction centred on the RSC. We computed mean logarithmic spectra, event-related spectral perturbations, topographic maps, and integrated dipoles, followed by a pre-clustering step in which each component was represented as a 10-dimensional vector with weights of 1 for mean logarithmic spectrum, 3 for event-related spectral perturbations, 1 for topographic maps, and 5 for dipole localization. Clustering was then performed with 5000 iterations to ensure reproducibility. We set the number of clusters to 15 and the outlier detection threshold to 3 standard deviations.

The choice of the number of clusters was based on the literature (Delaux et al., 2021), which suggested that having fewer clusters than components per subject yields better results. This clustering step, developed within the BeMobil processing pipeline, allows the addition of a region of interest. In our case, this was the RSC with MNI coordinates: x = 0, y = -55, z = 15. The choice of coordinates was based on the comparisons proposed by Delaux et al. (2021), and given the assumed absence of lateralization, we set x to 0. After 5000 clustering iterations, each clustering solution was evaluated based on six metrics:

1. Number of participants represented in the cluster (each cluster must contain at least one IC per subject, weight = 8).
2. Number of components per subject (each cluster should maximize ICs/participant, weight = -3).
3. Cluster dispersion (normalized by the number of ICs in the cluster, weight = -1).
4. Mean residual variance (weight = -1).
5. Distance between the cluster centroid and the region of interest (in this case the RSC: 0, -55, 15, weight = -3).
6. Mahalanobis distance from the median of the clustering solutions (weight = -1).

The final retained solution includes 33 subjects, with a mean residual variance of 0.048 ± 0.031 and a mean distance from the ROI of 24.46 ± 9.66 mm. In the case of multiple ICs for the same subject, we decided to select the IC with the lowest residual variance.

### Statistical analyses

Analyses were performed on participants’ accuracy scores, measured as the absolute angular error in degrees, as a function of perspective angles and angle to compute for physical rotation values (*i.e.,* the angle between the imagined perspective and the target) (**Figure 1.C**). We excluded all trials in which participants did not rotate in the desired direction (*i.e.,* the shorter one, as specified in the task’s instructions), as well as outliers using the quantile range on individual participant error distributions, which allowed us to retain 89.31% of the total trials (corresponding to a mean of 423.83 valid trials per participant, with *SE* = 59.14).

We then performed statistical analyses using R statistical software packages (version 4.4.0; R Foundation for Statistical Computing, Vienna, Austria) with R studio (version 2024.09.1). More specifically, we fitted linear mixed models (LMMs) using the *lme4* package (Bates et al., 2015) as well as generalized additive mixed models (GAMMs) using the *mgcv* (Wood, 2025) and *nlme* packages (Pinheiro et al., 2025) to account for the complexity of the relationships between our variables. The models were fitted using Maximum Likelihood (ML) and compared with the Bayesian Information Criterion (BIC, Schwarz, 1978). The selected model was then refitted using Restricted Maximum Likelihood (REML). We evaluated and modelled residual heteroscedasticity using *varPower*. This enabled us to address the heteroscedasticity of the residuals, given that the variability of the results increases with increasing angles. Approximate *p-*values for both parametric and smooth variables were extracted from the output of *summary(modelX$gam)*. Estimated marginal means (EMMs) were calculated using the *emmeans* package (Lenth et al., 2025) and are the values reported hereafter. Normality of the residuals and homoscedasticity were assessed using quantile-quantile plots and box plots, respectively. To account for the circular nature of our data, in a follow-up analysis we extracted the signed error values and quantified the consistency of participant’s responses using the Mean Vector Length (MVL), a metric ranging between 0 (*i.e.,* randomly dispersed values around the circle), and 1 (*i.e.,* highly concentrated values). We also extracted the Mean Direction Angle (MDA), reflecting the central direction of participant’s responses.

For the EEG analyses, following the clustering procedure, we extracted the time-frequency representations of the clustered components between 2 and 40 Hz using the *superlet* approach, which is well-suited for detecting transient bursts of activity (Moca et al., 2021). We applied a baseline correction using the interval from –500 to –100 ms (Cohen, 2014) and performed single-trial z-score normalization based on this time window (Grandchamp & Delorme, 2011) to extract the event-related spectral perturbations (ERSPs). Cluster-based permutation testing was then conducted on the ERSPs, using a statistical threshold of *p* < .05 (Maris & Oostenveld, 2007).

## Results

### Behavioural Analysis

#### Evolution of mean error angle according to perspective taking and rotation angle

To test our initial hypothesis, that the size of the perspective shift would influence the mean absolute angular error differently compared to the rotation angle, we first fitted a series of LMMs. These models included perspective-taking and rotation angles as fixed factors, and participant as a random intercept, with and without interaction between the variables (**Supplementary 1**). Given the large sample size and the complexity of the relationships in our data, we selected the best-fitting model using the BIC. The selected LMM revealed both a significant main effect of perspective-taking (*F*_(8, 9846.4)_ = 16.22, *p <* .001, *η^²^* = 0.01, 95% CI = [0.01, 0.02]) and rotation angle (*F*_(5, 9846.7)_ = 139.57 *p <* .001, *η^²^* = 0.07, 95% CI = [0.06, 0.08]). All details of this analysis are available in **Supplementary 1.** However, because the relation between rotation angle and the mean absolute angular error was non-linear we also compared models that included rotation angle in non-linear forms using GAMMs. This approach allowed us to capture the complex relationships between our variables (see **Figure 2**), without imposing a particular shape *a priori* (Crawley, 2015). More specifically, the following model to predict the mean absolute angular error includes a parametric term to capture the linear effect of perspective-taking, a smooth term for rotation angle to capture its non-linear effects, and a random intercept for participant:

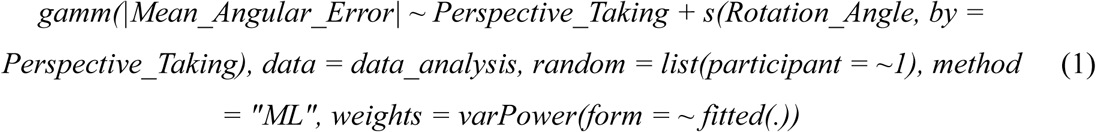

**Figure 2.**
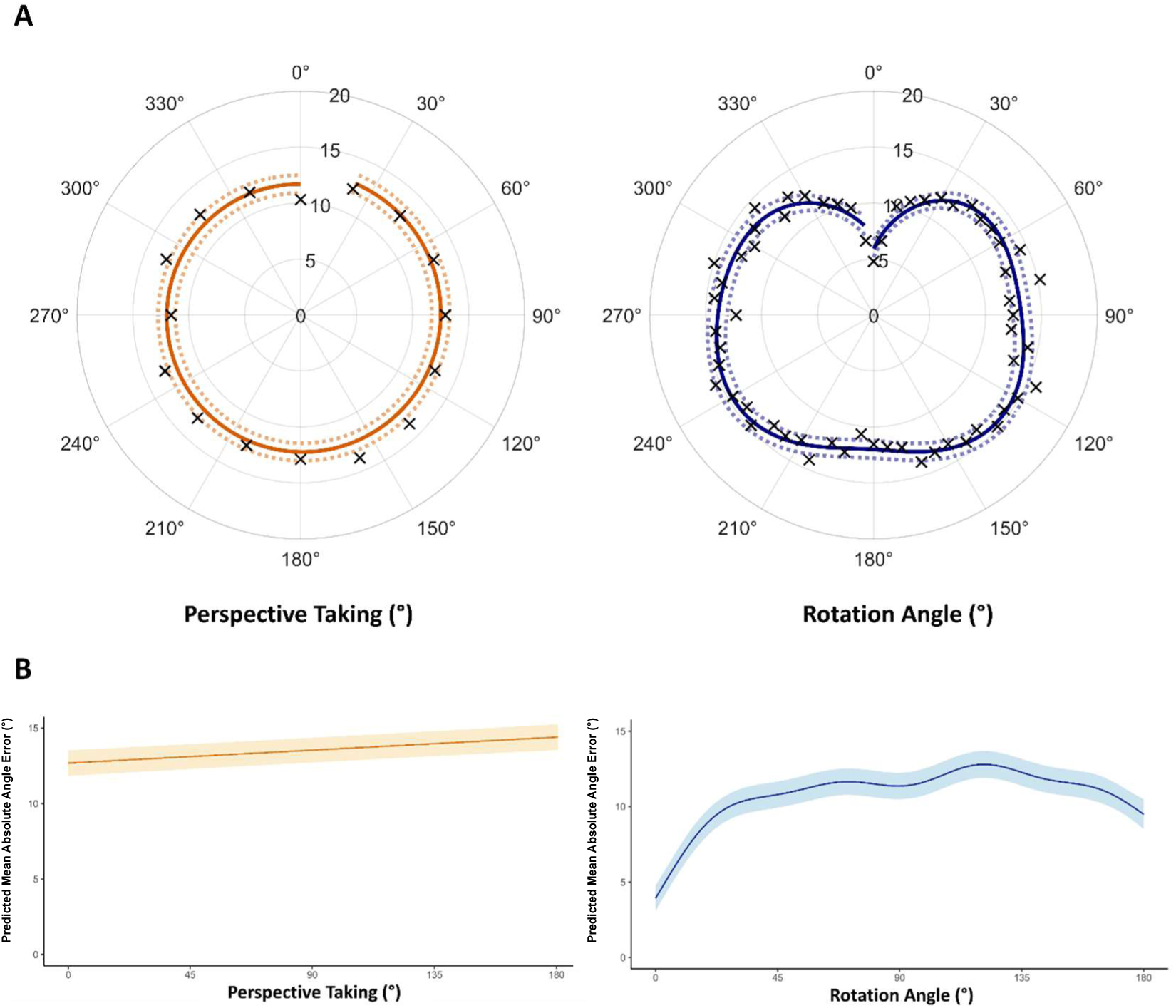
Two different patterns of angular error. **A.** Polar plots of the mean angular error (in degrees) of participants according to the perspective taking (left panel) and the rotation angle (right panel). the plots display the smoothed error lines (bold lines), and the 95% confidence intervals (shaded areas) based on the GAMM model including perspective taking as a fixed effect, rotation angle as a smooth predictor, and participant as a random effect. Mean angular errors are shown with black crosses, grouped by rotation angle in the left panel and by perspective taking in the right panel. A clear linear pattern is observed for angular errors according to perspective taking, in contrast to rotation angle, where errors follow a non-linear trend. **B.** These plots show the same data as in A, but with values corrected to lie in the range 0° to 180°. Mean angular errors are not displayed here. The same patterns as previously are highlighted, and we used these corrected angles for all the analyses.

**Figure 2** illustrates predictions of the best-fitting GAMM, suggesting a linear effect of perspective taking and a nonlinear effect of rotation angle (**Figure 2.A**), indicating distinct contributions of each variable to participants’ performance. These patterns were consistent across both sides of rotation and of perspective (**Figure 2.B**), confirming the absence of difference between left and right rotation as mentioned previously.

The results also showed a specific pattern at 0° perspective taking, suggesting a specific process in this condition, which involves only a physical rotation without a change in viewpoint. To address this possibility, we compared interaction and no-interaction GAMMs models on a subset of data excluding the 0° perspective condition (all the details of the GAMM are presented in **Supplementary 2**). Considering the BIC, the model without interaction fitted the best, suggesting that previously observed interaction between perspective taking and rotation angle was indeed primarily driven by this condition, and that the effect of rotation angle on error does not vary significantly across the other perspective angles.

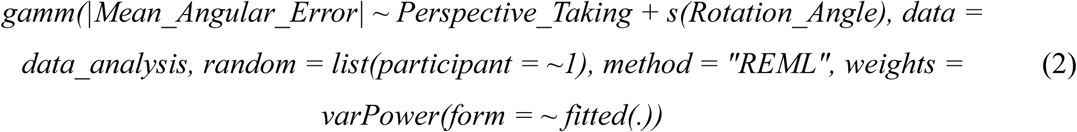

The final selected GAMM explained 35% of the variance when considering only the effect of fixed predictors (marginal R² = .35). We observed a significant parametric effect of perspective-taking (reference = 22.5°) and specifically for perspective angles of 67.5° (*β =* 0.56, 95% CI [0.02, 1.16], *p* = .044), 112.5° (*β =* 0.66, 95% CI [0.08, 1.24], *p* = .026), 135° (*β =* 0.70, 95% CI [0.12, 1.28], *p* = .017), and 157.5° (*β =* 0.78, 95% CI [0.20, 1.36], *p* = .009), indicating that these angles led to significantly greater mean absolute angular error. However, the results obtained from perspectives at 45°, 90°, and 180° did not differ from the 22.5° reference (all *p > .16*). The results are in line with our initial hypothesis, showing a linear increase in error with perspective-taking angle, consistent with a linear pattern.

A significant smooth term for rotation angle remained, with effective degrees of freedom deviating from 1 *(edf* = 7.79, *approx. p <* .001), confirming a complex and non-linear relationship between rotation angle and the mean absolute angular error, but not modulated by perspective-taking. Previous analyses did not reveal any interaction between perspective-taking and rotation angle. The latter showed a complex relationship with angular error and was probably influenced by execution error. As our experimental design did not allow us to verify this hypothesis, we decided to remove rotation angle from the subsequent models.

#### The influence of congruency and perspective taking on the angular error

We then investigated the effect of the congruency between the imagined perspective and the angle to be computed for the physical rotation. Incongruent trials are thought to require greater cognitive demands (*e.g.*, conflict resolution), which should lead to lower accuracy compared to congruent trials. Given that we demonstrated a linear relationship between the mean absolute angular error and the perspective taking and that GAMM is designed to capture non-linear effects, we decided to use LMM for follow-up analysis, to avoid introduce unnecessary complexity. Accordingly, we used the Akaike Information Criterion (AIC) for model selection, as model complexity was less of a concern in this context. Based on the lowest AIC value (**Supplementary 3**), we selected the following model:

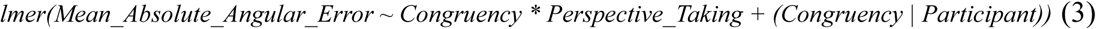

The results highlighted a main effect of congruency (*F*_(1, 37)_ = 7.50, *p =* .008, *η^²^_p_* = 0.18, 95% CI = [0.01, 0.39]) with participants showing better performance in the congruent condition (*EMM =* 12.80; *SE =* 0.41) than in the incongruent one (*EMM =* 13.70; *SE =* 0.45). We also reported a main effect of perspective-taking (*F*_(6, 444)_ = 4.41, *p <* .001, *η^²^_p_* = 0.06, 95% CI = [0.01, 0.09]) as well as a significant main interaction between congruency and perspective-taking (*F*_(6, 444)_ = 2.81, *p =* .01, *η^²^_p_* = 0.04, 95% CI = [0.00, 0.07]). Post hoc analysis of the interaction between congruency and perspective taking revealed that the congruency effect was only present for values of perspective taking above 90° (**Figure 3.A**). At these larger angles of 90°, 112.5°, and 135°, the congruency effect was statistically significant: 90° (*t*_(240)_ = -2.75, *p =* .006, *d =* -0.72, 95% CI [-1.24, -0.20]), 112.5° (*t*_(240)_ = -3.32, *p =* .001, *d =* -0.87, 95% CI [-1.40, -0.35]), and 135° (*t*_(240)_ = -3.13, *p =* .002, *d =* -0.82, 95% CI [-1.34, -0.30]), with incongruent trials showing significantly lower precision (*i.e.,* higher angular errors) compared to congruent ones. In contrast, no significant congruency effect was found for lower values of perspective taking (all *p >* .122), except for the perspective at 157.5°. This result suggested that participants made a greater error when both representations were incongruent, but only when they had to imagine a large shift in perspective.

**Figure 3.**
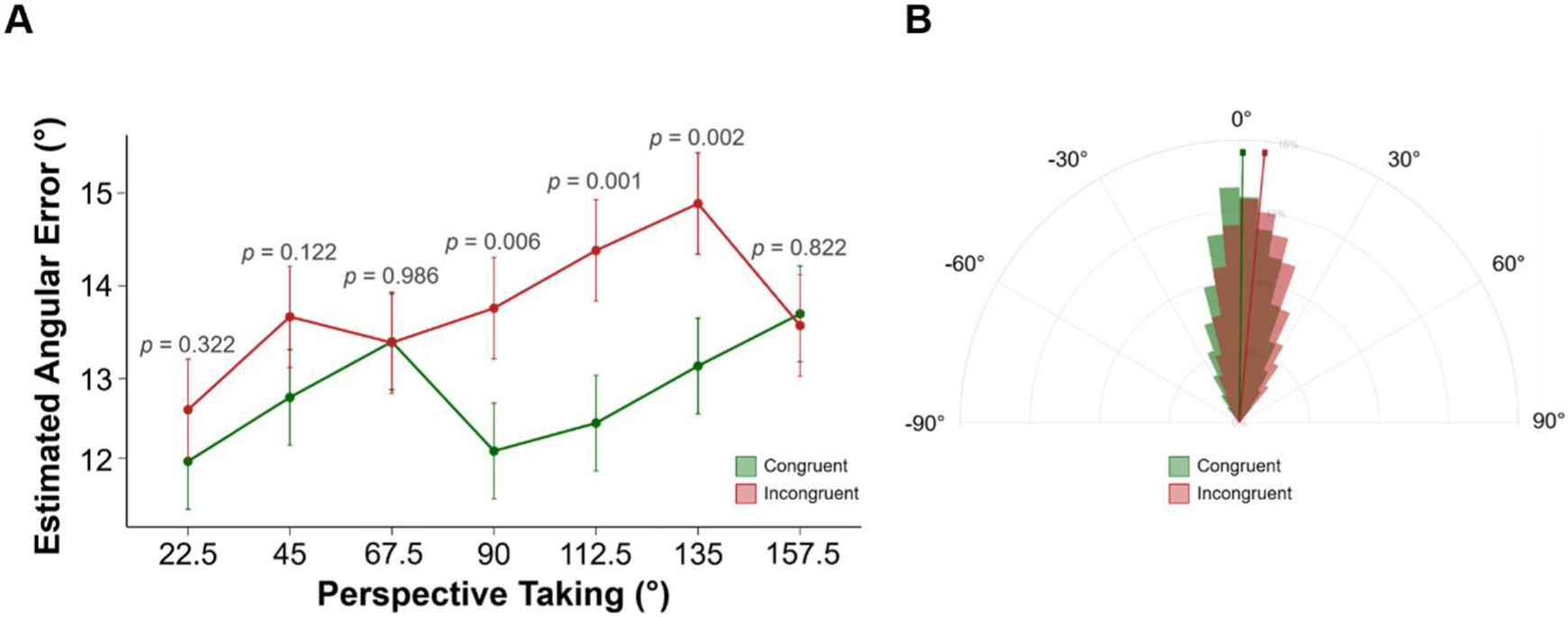
**A.** Graphical representation of the linear mixed-effects model showing the interaction between congruency and perspective taking on the estimated marginal means of angular error (± standard error). **B.** Polar histogram showing the proportion of error types (overshoot vs. undershoot) in the congruent and incongruent conditions. The length of the bold arrow represents the Mean Vector Length (MVL), indicating response consistency, and its direction corresponds to the Mean Direction Angle.

Interestingly, in a follow-up analysis we also observed an effect of the type of error between the two conditions. Participants tended to rotate more than needed (referred to as overshoot) in the incongruent condition (*EMM =* 63.0%; *SE =* 2.97%) compared to the congruent condition (*EMM =* 51.2%; *SE =* 2.79%; *F*_(1, 37)_ = 63.44, *p <* .001, *η^²^_p_* = 0.63, 95% CI = [0.43, 0.75]). There was no effect of the size of the perspective-taking angle (*p =* .32), suggesting that the effect relied solely on the congruency effect, participants performing a larger than required rotation in the incongruent condition only. As showed in **Figure 3.B** response consistency, indicated by the MVL, varied with perspective taking (*F*_(6, 481)_ = 4.24, *p <* .001, *η^²^_p_* = 0.05, 95% CI = [0.01, 0.08]), but did not differ by congruency (*F*_(1, 481)_ = 0.39, *p =* .533): with no difference between congruent trials (*MVL =* 0.969, *SE =* 0.001) and incongruent trials (*MVL =* 0.970, *SE =* 0.001). The interaction was not significant between the two variables (*F*_(6, 481)_ = 0.77, *p =* .590). In contrast, congruency shifted the mean direction angle (MDA; *F*_(1, 481)_ = 206.81, *p <* .001, *η^²^_p_* = 0.30, 95% CI = [0.24, 0.36], with larger MDAs in the incongruent condition (*EMM =* 5.24°; *SE =* 1.11) than the congruent condition (*EMM =* 0.49°; *SE =* 1.11). There was no main effect of perspective taking on MDA (*F*_(6, 481)_ = 0.69, *p =* .660), but the interaction between congruency and perspective taking was significant (*F*_(6, 481)_ = 6.800, *p <* .001, *η^²^_p_* = 0.078, 95% CI = [0.03, 0.12]). Post hoc analysis showed a reliable congruency effect at every perspective taking value (all *p <*.05). Importantly, the effect size (Cohen’s d) increased from small (*d =* -0.67 at 22.5°) to very large (*d =* -2.11 at 112.5°), before decreasing (*d =* -0.57° at 157.5°). Full details of the post hoc comparisons are provided in **Supplementary 4**. These results suggest that incongruency increases the general tendency toward overshoot without broadening the dispersion of the responses, and that this bias becomes stronger as the perspective-taking angle increase.

### iVTT performance correlates with perspective-taking value but only for congruent trials

We performed a Pearson’s correlation analysis on participants’ performance in the PTT and iVTT tasks, splitting the data into congruent and incongruent trials. Analyses were performed on data from 36 participants as two were excluded because they misunderstood the PTT’s task.

First, we replicated the main results from He et al. (2022), confirming a correlation between the absolute angular error for the iVTT and the PTT (*r =* .41, *p =* .01). We then further explored this relationship by integrating the congruency condition (*i.e.,* separating congruent and incongruent trials for the iVTT but not for the PTT as there were not enough balanced trials). Our analysis revealed that the correlation with performance for the iVTT and the PTT was only present for the congruent iVTT condition (*r =* .47, *t*_(34)_ = 3.13, *p =* .004, 95% CI [0.17, 0.69]), while incongruent trials did not correlate significantly with the error in the PTT (*r =* .24, *t*_(34)_ = 1.46, *p =* .15, 95% CI [-0.09, 0.53]) (**Figure 4**). This result suggests that visuo-spatial skills were not the only ones at play in the incongruent condition and that other factors such as the interaction between cognitive processes and/or motor preparation could also impact the performances.

**Figure 4.**
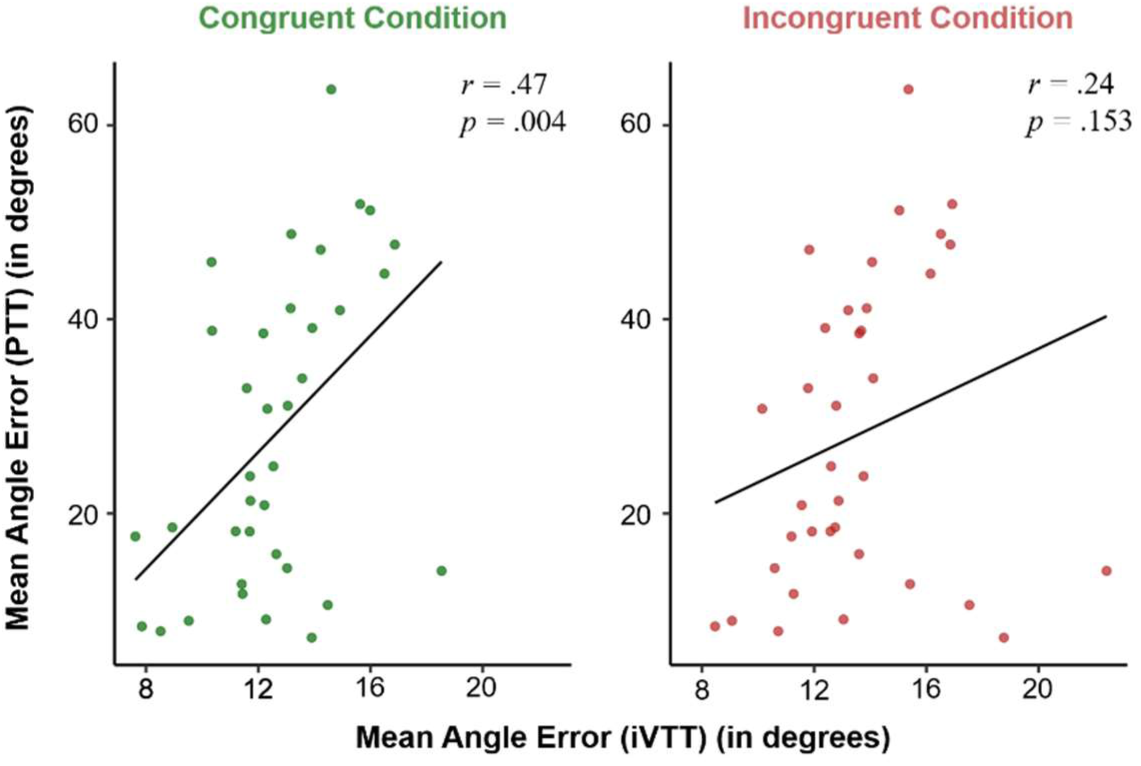
Correlation between performances for the iVTT and the PTT (distinguishing between congruent [left panel] and incongruent [right panel] trial), expressed in degrees. Each point represents a participant, and the linear regression line is shown in black.

### EEG Results

For the EEG data, we first focussed our analysis on a cluster of dipoles reconstructed around the RSC. This was motivated by recent evidence suggesting the RSC is a hub for spatial cognition, that is proposed to be involved in perspective taking despite few neural results (Alexander et al., 2023). Furthermore, in previous studies the RSC has been consistently implicated in tasks involving active rotation (Gramann et al., 2021; Naveilhan et al., 2025), making this region of particular interest to understand perspective-taking tasks that include physical rotation.

Firstly, we investigated the neural dynamics upon presentation of the map, that is, when participants were tasked to imagine taking the perspective of the arrow and computing the vector toward the blue dot. We performed analyses on the activity reconstructed around the RSC regions of 31 participants (**Figure 5.A**). Results highlighted a significant main effect of perspective taking in the RSC region (**Figure 5.B**). Specifically, this effect was driven by a cluster between 480ms and 740ms, restricted to frequencies between 10 and 17Hz, corresponding to high-alpha- and low-beta-band activity. Then, we performed a similar analysis, but using the angle to compute for physical rotation as the main independent variable (**Figure 5.C**). Here, results also showed a main effect of the angle to compute for physical rotation on RSC neural dynamics. However, this time the effect was driven by a later cluster (between 750 and 1150ms) at higher frequencies (between 22 and 28Hz), in the high-beta band. Thus, these results indicate a sequential process associated with perspective taking and angle computation. Overall, our findings suggest that both tasks are underpinned by activity in the RSC region; however, the first is associated with alpha-beta activity, whereas the second is linked to high-beta activity.

**Figure 5.**
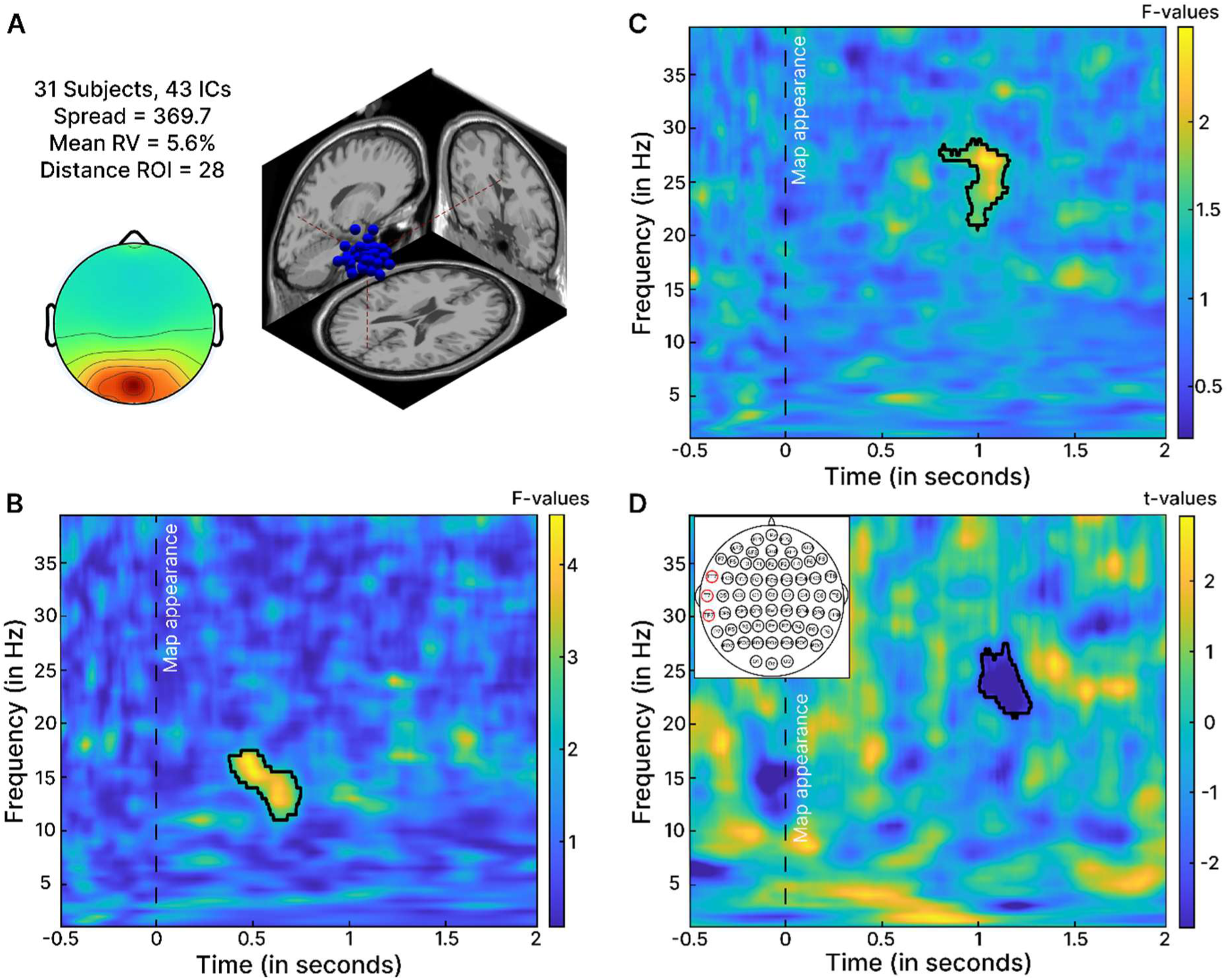
Presentation of the EEG results for map presentation phase, lasting 2 seconds. A. Cluster of dipoles reconstructed around the RSC. The best clustering solution included 43 dipoles, from 31 subjects, we then selected the one that had the lowest residual variance and was closest to the region of interest. B. Results of cluster-based permutation testing of time-frequency decomposition for the perspective taking (black lines indicate corrected p-values < .05). C. Results of cluster-based permutation testing of time-frequency decomposition for the angle to compute for physical rotation. D. As we reported no main effect of the congruency in the cluster of RSC dipoles, we then looked at clusters of electrodes. The figure presents the only cluster of electrodes with significant effect of the congruency, a cluster including electrodes FT7/T7/TP7, encompassing the lTPJ. The black line indicates cluster based corrected p-values below the .05 threshold adjusted with Bonferroni correction for the number of cluster (so *pcluster* < .005), negative *t-values* indicate less activity for the congruent condition.

Finally, we performed a similar analysis to investigate the effect of congruency between the perspective and the angle to compute for physical rotation. Results reported no differences in the RSC, even when using less conservative corrections for multiple comparisons. We therefore decided to extend our analyses to other regions in an exploratory way, and performed analyses at the electrode level, clustering the electrodes around functional brain regions as proposed in previous work (Harel et al., 2016; Kranczioch et al., 2006). This led to 16 clusters of electrodes, from which we extracted time-frequency activity and performed cluster-based permutation testing to assess the effect of the congruency (**Figure 5.D**). Only the cluster spanning electrodes FT7/T7/TP7 in the left temporo-parietal region survived after cluster-based correction with a Bonferroni adjusted *p-threshold* of 0.003 (0.05 / 16 clusters of electrodes). In these electrodes the effect of the congruency was driven by a cluster of activity between 990 and 1280ms at 21 and 27 Hz, thus in the high-beta-band frequencies with higher activity for the incongruent condition.

## Discussion

The aim of this study was to investigate the neurocognitive processes involved in spatial perspective taking and the execution of physical rotation using the immersive Viewpoint Transformation Task (iVTT). The fine-grained modulation across many perspective and rotation angle combinations revealed a different pattern for the two processes. While we observed a slight linear decrease in performance with increasing perspective shift, a non-linear relationship has been reported between performance and the angle required for physical rotation. These distinct patterns of angular error were mirrored in the EEG results around the retrosplenial complex region. Specifically, perspective-taking was associated with alpha-band activity, reflecting the mental shift in spatial reference frames required to perform the task, whereas the computation of the physical rotation angle emerged later and was predominantly reflected in beta-band activity. Finally, we observed an effect of congruency between the direction of perspective-taking and the physical rotation. When these directions were incongruent, performance declined, with participants tending to overshoot the required rotation. This congruency effect was also evident at the neural level, characterized by an increased beta-band activity in electrodes covering the left temporo-parietal junction (lTPJ).

### Dissociable behavioural patterns of perspective and rotation angle processing

In the first instance, our aim was to investigate the factors that contribute to the angular error of pointing during a perspective taking task involving physical rotation. In traditional perspective taking tests, participants are required to indicate on a circle the direction of a target object after taking a perspective different from their own, while in the present task participants had to physically rotate themselves to face this direction. Consistently with previous studies, we observed an increase in participants’ angular error when the size of the perspective angle increased (Kessler & Thomson, 2010; Kozhevnikov & Hegarty, 2001; Surtees et al., 2013). However, this effect was less pronounced than in the seminal work by Kozhevnikov & Hegarty (2001a). While they reported an increase of approximately 0.24° of error per degree of perspective, the increase was only 0.0093° in our study. Interestingly, the effect of the perspective shift was also smaller in the immersive study carried out by He et al. (2022), with our reanalysis indicating a beta estimate of 0.1072°. Thus, physical immersion in the environment may have reduced the effect of perspective taking on participants’ performance, possibly related to the embodied mechanisms afforded by real-time sensorimotor and proprioceptive alignment in virtual reality (Renata et al., 2024).

Our results also showed a similar pattern of linear increase between the error and the angle to compute. This relation was expected as during the rotation participants accumulate noisy velocity information over time leading to a noisier probability distribution of their heading (Harootonian et al., 2022). Chrastil & Warren (2017, 2021) interpreted this as an “execution error” corresponding to the discrepancy between the intended movement and the actual behavioural response. However, in our results the observed pattern was more complex and non-linear. Specifically, in addition to the expected linear increase of angular error up to 120°, we also showed that error then declined for angles closer to the antero-posterior axis (closer to 180°). This result highlights how spatial representations are organized around principal environmental and bodily axes (Kelly & McNamara, 2010; Newman et al., 2021; Rump & McNamara, 2013). This may be even more pronounced in virtual reality tasks in which self-motion cues markedly enhance these alignment effects by co-activating vestibular, proprioceptive, and visual signals, in order to minimize the updating variance when rotations align with bodily and environmental axes (Kelly et al., 2008; Riecke et al., 2015). On a related note, we also observed a slightly decreased error when the perspective angle matched the main body axes (0-180° for the antero-posterior and 90° left-right for the lateral axis) **(Supplementary 5**).

#### Alpha and beta activity in the RSC support perspective shift and physical rotation respectively

As expected, the effect of mental rotation of perspective emerged earlier in the temporal sequence compared to the angle computation for physical rotation, consistent with our experimental design. Moreover, time-frequency analyses revealed a cluster of activity in the alpha-band in the RSC, interpreted as a mental projection of oneself into the indicated location and orientation on the map. This finding is consistent with a previous EEG study from Gramann et al., (2010), who showed that modulation of alpha-band activity in the RSC preceded heading changes in a navigation task, reflecting the transformation of reference frames. Later work extended this result, reporting that the magnitude of RSC alpha modulation correlated with task accuracy, further underscoring its role as a key mechanism for reference frame transformation (Chiu et al., 2012; Lin et al., 2015). Our RSC findings are consistent with previous fMRI studies highlighting the RSC’s role in imagined perspective shifts, notably during the anticipation of viewpoint changes (Sulpizio et al., 2013) or the integration of different spatial viewpoints (Alexander et al., 2023; Dhindsa et al., 2014; Gomez et al., 2014; Marchette et al., 2014).

We also observed a delayed modulation of beta activity in the RSC that scaled with the magnitude of the angle to compute for physical rotation. The involvement of the RSC in this later phase aligns with its established role in route planning and encoding of future goal locations (Brown et al., 2016; Savelli & Knierim, 2010; Vann et al., 2009). In the same way, animal studies have revealed projections from the RSC to the secondary motor cortex, suggesting a direct pathway through which spatial representations can be translated into motor plans (Bicanski & Burgess, 2016; Yamawaki et al., 2016). These findings resonate with the known functions of the posterior parietal cortex, which serves as an integrative hub for visual, somatosensory, and motor signals to perform sensorimotor transformations required for goal-directed movement (Andersen & Buneo, 2002; Freedman & Ibos, 2018; Kuang et al., 2016) and thus suggests motor precomputation processes within the RSC. Supporting this view, electrophysiological evidence has linked beta-band activity in the posterior parietal cortex to both executed and imagined movements (Brovelli et al., 2004; MacKay & Mendoncca 1995). Overall, our findings are consistent with the role of the RSC integrating both motor and non-motor aspects related to spatial orientation (Alexander et al., 2023).

#### Incongruency effect between mental perspective shift and physical rotation leads to lower performances and beta activity in the lTPJ

We reported an increased angular error for incongruent trials, characterized by a significant tendency of participants to overshoot the target when there was a mismatch between the imagined direction (*i.e.,* the perspective shift) and the physical rotation. This suggests that the additional complexity induced by incongruency may require greater cognitive resources. Interestingly, this incongruency effect on the angular error appeared only when the perspective shift was greater than 90°, consistent with previous findings from Kessler & Thomson (2010). These results could be interpreted in terms of the different strategies employed for small versus large perspective shifts. Participants presumably relied on allocentric mental rotation for smaller shifts and switched to an egocentric perspective-taking strategy when shifts were more substantial (Geva & Henik, 2019; Hegarty & Waller, 2004; Surtees et al., 2013). For larger perspective shifts (> 90°), perspective-taking is more likely to be used. In such cases, incongruency between the direction of the physical rotation to compute and the perspective taking may require increased top-down control to suppress the incorrect response and disregard the actual perspective which can interfere with the intended one (Keehner et al., 2006; Puls & May, 2020; Zwickel & Müller, 2013). This interpretation is strengthened by the tendency of participants to overshoot the rotation in the incongruent condition, with a linear increase for perspective bigger than 90°. Thus, the incongruent perspective may be interpreted as vector sum, significantly impacting the physical rotation phase for larger perspective shift (Segen et al., 2021).

This conflict between spatial representations reported at the behavioural level was also present at the neural level, with differences between congruent and incongruent trials in the brain activity in electrodes encompassing the lTPJ. The involvement of this region in our study is consistent with previous work highlighting the central role of this brain region when signal prediction errors are used to update internal models when the anticipated and the required angle to perform diverge (Doricchi et al., 2022). The observed increase in beta-band activity over these electrodes may mark moments of such updating, reflecting the transient inhibition and reshaping of sensory information before the correct motor response is initiated (Lundqvist et al., 2024). In line with this interpretation, Spitzer & Haegens (2017) proposed that brief beta bursts initiate top-down inhibitory control by temporarily suppressing competing neural ensembles to facilitate the selective reactivation of task-relevant content (Haegens et al., 2017; Limanowski et al., 2020). In our study, the observed increase in beta-band activity may be interpreted as the detection of a mismatch between the imagined perspective shift and the required direction of rotation, which is used to update the direction primed during perspective taking.

## Conclusion

In conclusion, this mobile EEG study allowed us to investigate the distinct neurocognitive processes underlying perspective-taking and physical rotation within an immersive spatial perspective taking task. Behaviourally, two distinct patterns of angular error emerged: a linear increase with the magnitude of imagined perspective shift, and a complex, non-linear pattern for physical rotation, with improved accuracy along the body’s main axis. At neural level, we reported that alpha-band activity in the RSC supports the transformation of spatial reference frames during imagined perspective shifts, while subsequent beta-band activity reflects motor precomputation of the required angular rotation. Importantly, incongruency between the imagined and executed directions leads to increased angular error and enhanced beta activity in electrodes encompassing the lTPJ, consistent with prediction error signalling and the updating of internal models. These findings highlight the sequential and dissociable dynamics of embodied spatial transformations and emphasize the value of immersive virtual environments for investigating the neural correlates of real-world navigation mechanisms. These results pave the way for further investigation in older adults, given that declines in spatial abilities have been shown to occur early in pathological aging, such as Alzheimer’s disease.

## Acknowledgements

This work was supported by the French government through the UCA^JEDI^ project managed by the National Research Agency (ANR-15-IDEX-01) and, in particular, by the interdisciplinary Institute for Modeling in Neuroscience and Cognition (NeuroMod) of Université Côte d’Azur.

## Code and Data Availability Statement

The complete set of raw data, codes, and stimuli generated for the analyses presented in this paper can be accessed through the OSF repository at: https://osf.io/7bzua/?view_only=c087063470cb4cd18682ae83767d339f

## Author Contributions

**Saulay-Carret Maud:** Methodology, Formal analysis, Investigation, Writing – Original Draft; **Naveilhan Clémen**t: Conceptualization, Methodology, Formal analysis, Investigation, Writing – Original Draft; **Xavier Corveleyn:** Writing – Review & Editing; **Stephen Ramanoël:** Conceptualization, Supervision, Writing – Review & Editing, Funding Acquisition, Project administration.

## Disclosure statement

No author involved with this research had any conflicts of interest. This work was approved by the local Ethical Committee of Université Côte d’Azur (CERNI-UCA authorization no. 2023-20) and participants provided informed consent before starting the experiment.

## Supplementary materials

### Supplementary 1: Comparison of LMMs using Bayesian Information Criterion

|  | Model formula (*lmer* function) | BIC |
| --- | --- | --- |
| **M_null** | *Angular_Error* ~ (1\|*Participant*) | 69313.59 |
| **M0** | *Angular_Error* ~ *Perspective_Taking* + (1\|*Participant*) | 69269.82 |
| **M0b** | *Angular_Error* ~ *Rotation_Angle* + (1\|*Participant*) | 69147.58 |
| **M1** | *Angular_Error* ~ *Perspective_Taking* + *Rotation_Angle* + (1\|*Participant*) | 69101.85 |
| **M2** | *Angular_Error* ~ *Perspective_Taking* * *Rotation_Angle* + (1\|*Participant*) | 69211.75 |
| **M3** | *Angular_Error* ~ *Perspective_Taking* + *ns*(*Rotation_Angle*,3) + (1\|*Participant*) | 68726.71 |
| **M4** | *Angular_Error* ~ *Perspective_Taking* * *ns*(*Rotation_Angle*,3) + (1\|*Participant*) | 68800.53 |
| **M5** | *Angular_Error* ~ *Perspective_Taking* + *ns*(*Rotation_Angle*,4) + (1\|*Participant*) | 68651.50 |
| **M6** | *Angular_Error* ~ *Perspective_Taking* * *ns*(*Rotation_Angle*,4) + (1\|*Participant*) | 68762.08 |
| **M7** | *Angular_Error* ~ *Perspective_Taking* + *ns*(*Rotation_Angle*,5) + (1\|*Participant*) | 68641.03 |
| **M8** | *Angular_Error* ~ *Perspective_Taking* * *ns*(*Rotation_Angle*,5) + (1\|*Participant*) | 68784.54 |
| **M9** | *Angular_Error* ~ *Perspective_Taking* + *ns*(*Rotation_Angle*,6) + (1\|*Participant*) | 68641.27 |
| **M10** | *Angular_Error* ~ *Perspective_Taking* * *ns*(*Rotation_Angle*,6) + (1\|*Participant*) | 68826.42 |

Note: *Angular\_Error* refers to the *Mean\_Absolute\_Angular\_Error*, as defined in the manuscript.

We compared twelve candidate models (M0-M10) using the Bayesian Information Criterion (BIC) to deal with both model fit and complexity. Based on BIC, Model M7 was selected for further analysis. We observed a significant main effect of perspective taking (*F*_(8, 9846.4)_ = 16.216, *p <* .001, *η^²^_p_* = 0.01, 95% CI = [0.01, 0.02]). Post hoc analysis showed an increase of the mean absolute angular error with perspective taking, peaking at 135° (*EMM* = 14.3; *SE =* 0.454), before slightly decreasing up to 180° (*EMM* = 13.8, *SE =* 0.463). An exception was observed at a perspective shift of 90° (*EMM* = 13.3, *SE =* 0.454). To capture the potential non-linear relationship between mean absolute angular error and rotation angle, we used natural splines with five degrees of freedom. We observed an overall significant main effect of rotation angle (*F*_(5, 9846.7)_ = 139.565, *p <* .001, *η^²^_p_ =* 0.066, 95% CI = [0.06, 0.08]). Four out of the five basis functions contributed significantly (respectively: *β*_1_ = 6.45, *SE_1_ =* 0.42; *β*_2_ = 9.11, *SE_2_ =* 0.52; *β*_3_ = 5.83, *SE_3_ =* 0.41; *β*_4_ = 14.73, *SE_4_ =* 0.77; with all *p <* .001), while the fifth one was not significant (*β*_5_ = 0.51, *SE_5_ =* 0.35, *t* = 1.49, *p =* .137). These results indicated a non-linear relationship between rotation angle and participant angular error, that was primarily explained by the first four spline components.

### Supplementary 2: Detailed results of GAMM (without interaction)

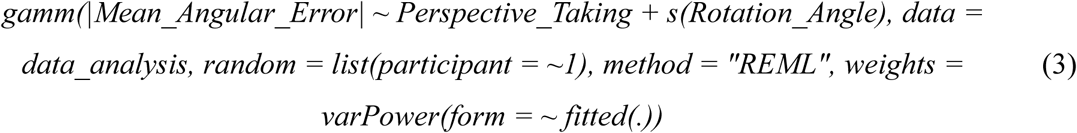

| Fixed Effects (reference = perspective of 22.5°) | $\beta$ (95% CI) | p-value |
| --- | --- | --- |
| Perspective-Taking of 45° | 0.26 (-0.3, 0.83) | .364 |
| Perspective-Taking of 67.5° | 0.59 (0.02, 1.16) | .044 |
| Perspective-Taking of 90° | -0.14 (-0.69, 0.42) | .630 |
| Perspective-Taking of 112.5° | 0.66 (0.08, 1.24) | .026 |
| Perspective-Taking of 135° | 0.70 (0.12, 1.28) | .018 |
| Perspective-Taking of 157.5° | 0.78 (0.20, 1.36) | .009 |
| Perspective-Taking of 180° | 0.42 (-0.17, 1.02) | .161 |
| Smooth Effects | Edf | p-value |
| Rotation Angle | 7.79 | < .001 |
| Random Effects |  |  |
| ICC | 0.765 |  |
| Marginal $R^2$/Conditional $R^2$ | 0.350/0.979 | |

Results from GAMMs including participant as random intercept, rotation angle as smooth term and perspective taking as parametric term; without interaction between those variables. We present *β estimates*, which corresponds to the estimate of the linear effect size of the predictor on the measured variable. For example, if we look at perspective angle 112.5°, we expect an increase of 0.66° in the mean absolute angular error for this value, compared to the reference value (22.5°). Regarding the smooth term, the *Edf* (effective degrees of freedom) indicates the non-linearity of the effect: the more *Edf* differs from 1, the more complex is the relationship between the corresponding variables; an *Edf* equal to 0 means no effect. The ICC (Intraclass Correlation Coefficient) quantifies the proportion of the residual variance (*i.e.,* variance not explained by the fixed effects) in the outcome that can be explained by random effects; it differs from the *ΔR²_conditinal_ – R²_marginal_*, which quantifies the additional variance explained by those random effects to the fixed-effects model.

### Supplementary 3: Comparison of LMMs included perspective taking and congruency, using Akaike Information Criterion

|  | Model formula (*lmer* function) | AIC |
| --- | --- | --- |
| **M0** | *Angular_Error* ~ (1\|Participant) | 2562.87 |
| **M1** | *Angular_Error* ~ (1\|Congruency) | 2800.80 |
| **M2** | *Angular_Error* ~ (Congruency\|Participant) | 2527.55 |
| **M3** | *Angular_Error* ~ Perspective_Taking + Congruency + (1\|Participant) | 2539.09 |
| **M4** | *Angular_Error* ~ Perspective_Taking * Congruency + (1\|Participant) | 2531.20 |
| **M5** | *Angular_Error* ~ Perspective_Taking + Congruency + (Congruency\|Participant) | 2512.85 |
| **M6** | *Angular_Error* ~ Perspective_Taking * Congruency + (Congruency\| Participant) | 2503.54 |

Note: Angular\_Error refers to the Mean\_Absolute\_Angular\_Error, as defined in the manuscript.

### Supplementary 4: Detailed post hoc results for the interaction between congruency and perspective taking on the MDA

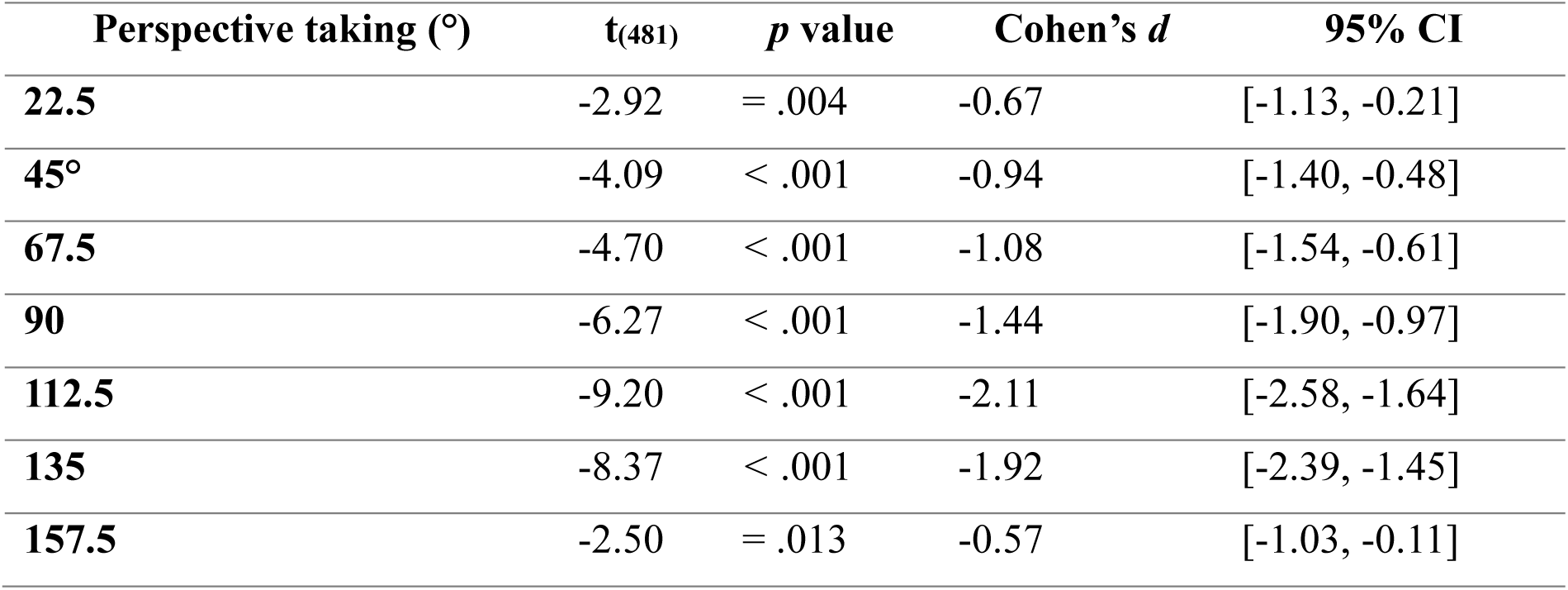

**Supplementary figure 1.**
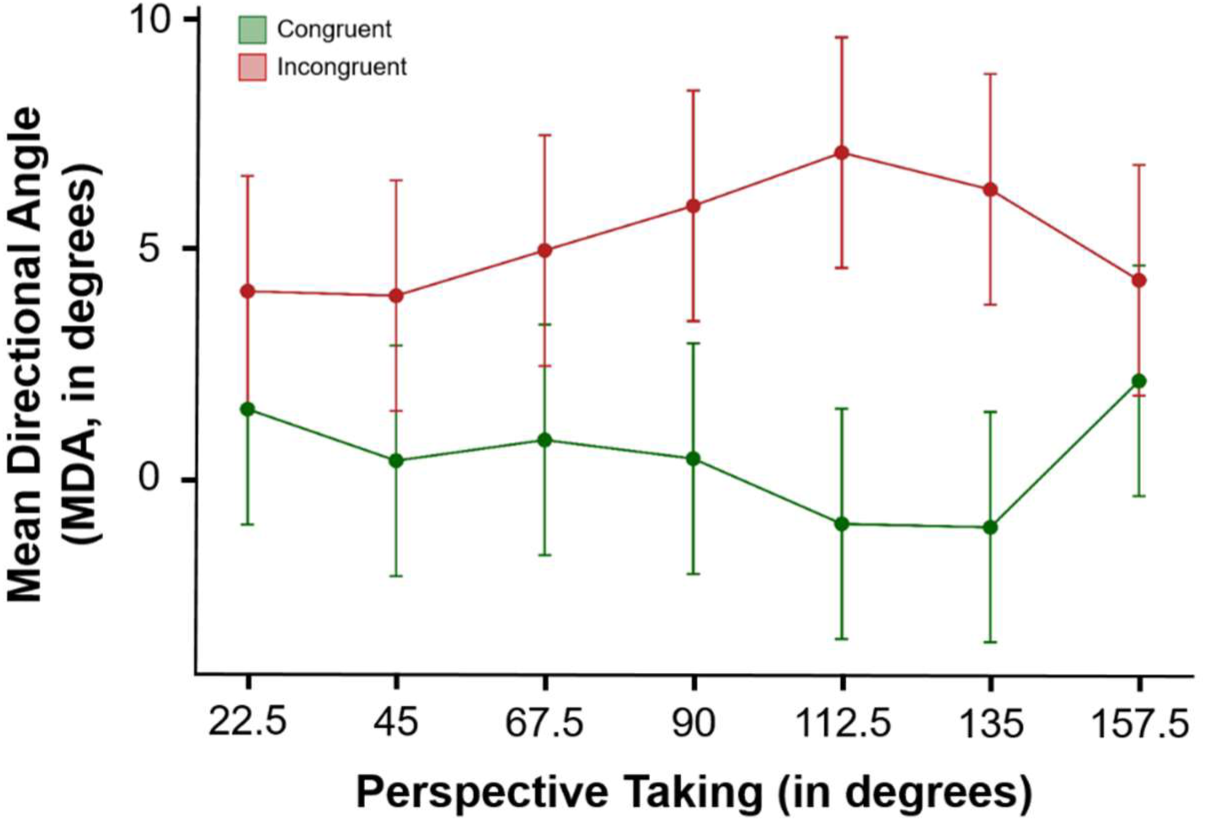
Estimated marginal means of Mean Direction Angle (MDA, in degrees; ± standard error) as a function of perspective-taking value (in degrees), for congruent (green) and incongruent (red) conditions.

### Supplementary 5: Predicted mean angular error (in degrees), based on a GAMM with interaction between perspective taking and rotation angle

**Supplementary figure 2.**
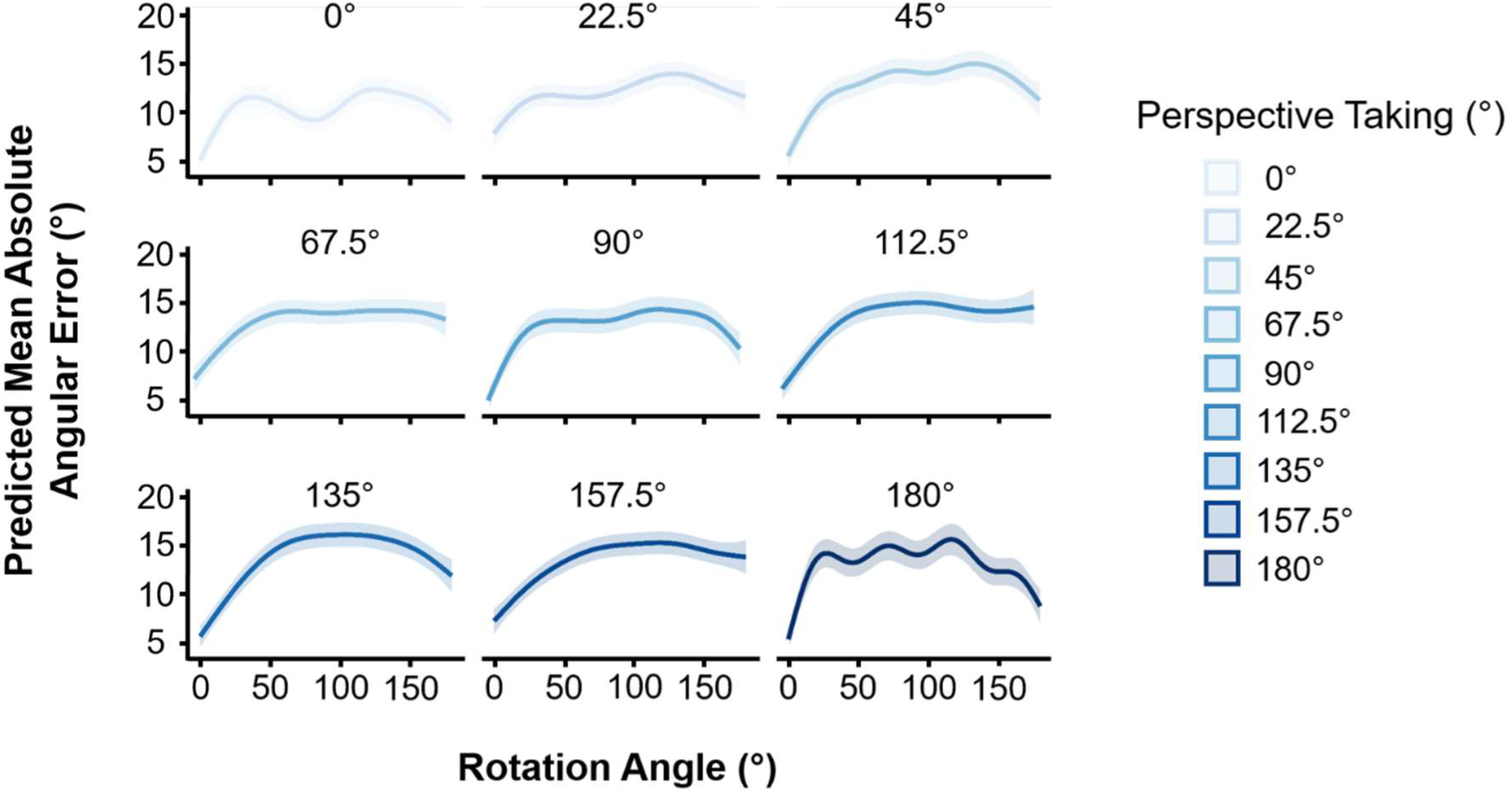
The plots display the predict mean angular error (in degrees), based on a GAMM including a smooth term for rotation angle varying with perspective taking, and a fixed effect of perspective taking. Each panel displays the smoothed error lines (bold lines), and the 95% confidence intervals (shaded areas) for one perspective condition (0°, 90°, 180°). This representation highlights the complex and non-linear relationship between rotation angle and angular error and shows reduced error when rotation angles align with body axes.

## Notes

https://osf.io/7bzua/?view_only=c087063470cb4cd18682ae83767d339f

### Competing Interest Statement

The authors have declared no competing interest.

